# *E. coli* OxyS non-coding RNA does not trigger RNAi in *C. elegans*

**DOI:** 10.1101/013029

**Authors:** Alper Akay, Peter Sarkies, Eric A. Miska

## Abstract

The discovery of RNA interference (RNAi) in *C. elegans* has had a major impact on scientific research, led to the rapid development of RNAi tools and has inspired RNA-based therapeutics. Astonishingly, nematodes, planaria and many insects take up double-stranded RNA (dsRNA) from their environment to elicit RNAi; the biological function of this mechanism is unclear. Recently, the *E. coli* OxyS non-coding RNA was shown to regulate gene expression in *C. elegans* when *E. coli* is offered as food. This was surprising given that *C. elegans* is unlikely to encounter *E. coli* in nature. To directly test the hypothesis that the *E. coli* OxyS non-coding RNA triggers the *C. elegans* RNAi pathway, we sequenced small RNAs from *C. elegans* after feeding with bacteria. We clearly demonstrate that the OxyS non-coding RNA does not trigger an RNAi response in *C. elegans*. We conclude that the biology of environmental RNAi remains to be discovered.

## Introduction

The biology underlying RNAi is still only partially understood. Bacteria engineered to express dsRNA homologous to *C. elegans* genes can induce RNAi when fed to the animals^1^; transgenic plants engineered to express dsRNA against genes of plant-parasitic nematodes kill these nematodes when used as hosts^2^. Why any animal, or indeed *C. elegans* in particular, should respond to environmental dsRNA, is an exciting question that remains unanswered. We recently discussed some potential roles of environmental RNAi in animal to animal communication (“social RNA”)^3^.

Environmental RNAi in *C. elegans* is a multi-step process. First, dsRNA molecules from food are taken up by intestinal cells, a process that requires the dsRNA transporter SID-2^4^. dsRNA and/or RNAi intermediates can move between cells and tissues of the animal, a phenomenon referred to as systemic RNAi, which requires SID-1 RNA channel and other protein factors^5,6^. Within cells dsRNA is processed by the DICER endonuclease (DCR-1)/RDE-4 complex, which generates short dsRNA^7–9^. Either of the strands of the short dsRNA, the 1° siRNA, is incorporated into the Argonaute protein RDE-1^10,11^ and finds its target through primary sequence complementarity (Watson-Crick base-pairing). Upon target recognition, the RDE-1/1° siRNA complex recruits RNA-dependent RNA polymerases (RdRPs) to generate 2° siRNAs, also known as 22G-RNAs^12^. 22G-RNAs associate with a different class of Argonaute proteins (WAGOs) and form the active RNAi effector complex. 22G-RNAs lead to gene silencing at both the post-transcriptional and transcriptional level^13,14^. Due to the RdRP amplification step, 22G-RNAs are highly abundant and are a key signature of RNAi in *C. elegans* (Supplementary Fig. 1).

**Figure 1:**
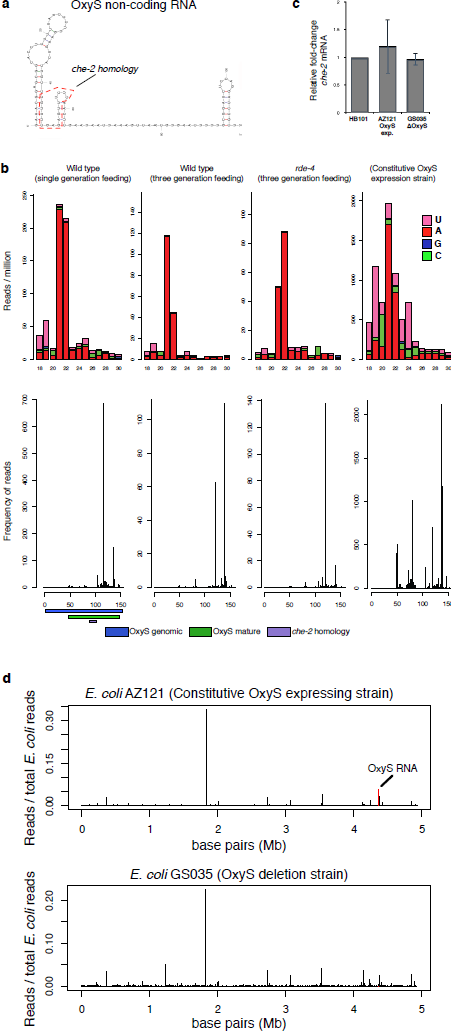
*E. coli* strain AZ121 with constitutive OxyS non-coding RNA expression does not induce RNAi in *C. elegans* upon feeding. **(a)** *E. coli* OxyS non-coding RNA structure. Dashed red lines highlight the possible homology region to *C. elegans che-2* gene. **(b)** Small RNAs from one or three generation fed *C. elegans* larvae and the small RNAs from *E. coli* AZ121 (constitutive OxyS expression strain) are mapped onto the OxyS RNA. Upper panels show the size and nucleotide distribution of mapped reads. Lower panels show the frequency of mapped reads along the OxyS non-coding RNA. **(c)** Relative fold-change of *che-2* mRNA in animals fed with either the constitutive OxyS expression strain (*E. coli* AZ121) or the ΔOxyS (*E. coli* GS035). Fold-change is calculated relative to the *che-2* mRNA levels in animals fed with the wild type *E. coli* strain HB101. **(d)** Small RNA sequencing reads from animals fed with either *E. coli* AZ121 (upper panel) or *E. coli* GS035 (lower panel) are mapped to the *E. coli* genome in windows of 5,000 bases. Red bar in the upper panel highlights the reads mapping to the OxyS transcript. There were no reads mapping to the OxyS transcript in the ΔOxyS GS035 strain (lower panel).

*C. elegans* is a common animal model and the standard food source in the laboratory is *E. coli*. In nature *C. elegans* is associated with soil and decaying organic matter and is likely a generalist feeding on microorganisms such as bacteria^15^. Curiously, the *E. coli* stress-induced non-coding RNA OxyS^16^ has a 17 nucleotides (nt) region complementary to the sequence of the *che-2* gene of *C. elegans*^17^ (Fig. 1a boxed area). Recently, Liu *et al.*^17^ reported that the OxyS non-coding RNA causes down-regulation of the *che-2* mRNA in *C. elegans* when expressed in the *E. coli* food. According to the genetic analysis of Liu *et al.*^17^, OxyS-dependent down-regulation of *che-2* requires some of the components of the RNAi and miRNA pathways in *C. elegans*. Specifically, *dcr-1* and *rde-4*, essential for long dsRNA processing during RNAi^8,9^, *rde-1*, the 1° siRNA binding Argonaute^11,18^, and *alg-1*, a microRNA (miRNA) pathway gene^10^, were all reported to be required for *che-2* mRNA regulation^17^. Surprisingly, the systemic RNAi genes sid-1^5^ and sid-2^4^ were found to be dispensible^17^. The authors concluded that the OxyS RNA elicits an RNAi response in *C. elegans* that results in gene expression changes and behavioral responses.

## Results

To test this hypothesis directly, we fed three different *E. coli* strains to *C. elegans* either for one or three generations: an OxyS over-expressing strain (AZ121), an OxyS deletion strain (GS035) and the common *C. elegans* food strain (HB101). We isolated total RNA from young adult stage animals and prepared 5’ independent small RNA libraries which enables the sequencing of both 1° and 2° siRNAs^12^ (Supplementary Fig. 1). We then mapped the small RNA reads against the OxyS non-coding RNA. We did not detect any small RNAs mapping to the OxyS RNA in animals fed with either the OxyS deletion strain or the HB101 *E. coli* strain (data not shown). In animals fed with the AZ121 strain, small RNAs mapping to the OxyS RNA show a peak at 21-22 nt both in one or three generation fed animals (Fig. 1b, top two panels). However, these small RNAs map exclusively to the 3’ region of the OxyS RNA. We did not detect any small RNAs mapping to the region of the OxyS RNA that is predicted to share sequence similarity to the *che-2* mRNA (Fig. 1b, bottom two panels). To test whether Dicer activity was involved in the generation of these small RNAs, we repeated the feeding experiment using *rde-4* mutant animals (Fig. 1b, top and bottom third panel). Small RNA reads from *rde-4* mutants showed a very similar pattern to those from wild-type animals with abundant 21-22 nt RNAs mapping exclusively to the 3’ region of the OxyS RNA. These results indicate that the DCR-1/RDE-4 complex is not required for the generation of these 21-22 nt small RNAs which originate from the OxyS RNA. Next, we checked for 22G-RNAs, which are the effectors of *C. elegans* RNAi pathway. We did not observe any 22G-RNAs mapping either against the OxyS RNA or to the *che-2* mRNA in our 5’ independent small RNA libraries, thus suggesting that there is no small RNA mediated silencing of *che-2*. Consistently, we also did not observe any down-regulation of the *che-2* mRNA upon AZ121 feeding (Fig. 1c).

Given the lack of engagement of the RNAi pathway, we speculated that the small RNAs mapping to the OxyS RNA might be generated by the bacteria and not by *C. elegans.* To test this we sequenced the small RNAs from the *E.coli* strains alone. In the AZ121 strain, we detected a very similar pattern of 21-22 nt small RNAs, mostly mapping to the 3’ end of the OxyS RNA and more abundant than the reads detected from the sequenced animals (Fig. 1b, rightmost panel). Thus small RNAs arising from the OxyS RNA are made within the bacteria and are not generated by the *C. elegans* RNAi pathway. We believe that these small RNAs are likely degradation products of the OxyS RNA and do not enter the RNAi pathway. We also mapped all the small RNAs to the *E. coli* genome to make sure that the OxyS RNA is expressed (Fig. 1d). In AZ121 strain, small RNA reads mapping to the OxyS RNA can be detected in abundance (Fig. 1d, top panel red line). In contrast, GS035 strain has no reads mapping to the OxyS RNA.

## Discussion

Here we show that small RNA sequencing can efficiently be used to elucidate possible regulatory roles of non-coding RNAs in *C. elegans*. Our analysis of the OxyS RNA as a trigger for RNAi in *C. elegans* clearly shows that OxyS non-coding RNA does not enter the canonical RNAi pathway in *C. elegans* upon feeding. We suggest that the previously described phenotypes in *C. elegans* upon OxyS non-coding RNA feeding^17^ are more likely to be caused by the differences within the bacterial strains. It is known that different bacteria can affect the physiology of the nematodes with respect to foraging, metabolism and ageing^19,20^. It remains possible that RNA produced by different bacteria in nature enters the *C. elegans* RNAi pathway; however, our results suggest that sequencing of the small RNA populations in *C. elegans* is essential to confirm this. We conclude that the biology of environmental RNAi remains to be discovered.

## Methods

*E. coli* HB101, AZ121 and GS035 strains were grown in LB media containing no antibiotics, ampicillin (100 μg/ml) or chloramphenicol (25 μg/ml) respectively (OD595=0.6-0.8). Cultures were centrifuged and the re-suspended pellets were seeded onto NGM plates. For one generation feeding experiments, animals were placed on the seeded NGM plates and allowed to grow until young adult stage. For three generation feeding experiments, L1 larval stage animals were placed on the seeded NGM plates and allowed to grow for three generations. Total RNA from harvested animals was isolated using the TriSure reagent (Bioline). Small RNA sequencing libraries of *C. elegans* were prepared using the NEBNext Multiplex Small RNA Library Prep set (NEB) and the small RNA sequencing libraries of *E. coli* are prepared using the TruSeq Small RNA set (Illumina). Libraries were sequenced on a MiSeq machine (Illumina). Data analysis was performed as previously described ^21^. qRT-PCR is performed with primers described in Liu et al. ^17^.

## Acknowledgements

We thank Gisela Storz for sharing *E. coli* AZ121 and GS035 strains. PS is funded by a research fellowship from the Gonville and Caius College, University of Cambridge. AA and EAM are supported by a Cancer Research UK programme grant to EAM.

**Competing financial interests:** The authors declare no competing financial interests.

**Author contributions:** AA initiated the project and performed the experiments. PS contributed to the experiments and analyzed the data. AA, PS and EAM developed the concepts and wrote the manuscript.

## References

1. Timmons, L., Court, D. L. & Fire, A. Ingestion of bacterially expressed dsRNAs can produce specific and potent genetic interference in Caenorhabditis elegans. Gene, 263, 103–112 (2001).

2. Yadav, B. C., Veluthambi, K. & Subramaniam, K. Host-generated double stranded RNA induces RNAi in plant-parasitic nematodes and protects the host from infection. Mol. Biochem. Parasitol. 148, 219–22 (2006).

3. Sarkies, P. & Miska, E. A. Molecular biology. Is there social RNA? Sci. (New York, NY) 341, 467–468 (2013).

4. Winston, W. M., Sutherlin, M., Wright, A. J., Feinberg, E. H. & Hunter, C. P. Caenorhabditis elegans SID-2 is required for environmental RNA interference. Proc. Natl. Acad. Sci. U. S. A. 104, 10565–10570 (2007).

5. Feinberg, E. H. & Hunter, C. P. Transport of dsRNA into cells by the transmembrane protein SID-1. Sci. (New York, NY) 301, 1545–1547 (2003).

6. Hinas, A., Wright, A. J. & Hunter, C. P. SID-5 is an endosome-associated protein required for efficient systemic RNAi in C. elegans. Curr. Biol. 22, 1938–1943 (2012).

7. Knight, S. W. & Bass, B. L. A role for the RNase III enzyme DCR-1 in RNA interference and germ line development in Caenorhabditis elegans. Sci. (New York, NY) 293, 2269–2271 (2001).

8. Parker, G. S., Eckert, D. M. & Bass, B. L. RDE-4 preferentially binds long dsRNA and its dimerization is necessary for cleavage of dsRNA to siRNA. RNA (New York, NY) 12, 807–818 (2006).

9. Tabara, H., Yigit, E., Siomi, H. & Mello, C. C. The dsRNA binding protein RDE-4 interacts with RDE-1, DCR-1, and a DExH-box helicase to direct RNAi in C. elegans. Cell 109, 861–871 (2002).

10. Grishok, A. et al. Genes and mechanisms related to RNA interference regulate expression of the small temporal RNAs that control C. elegans developmental timing. Cell 106, 23–34 (2001).

11. Yigit, E. et al. Analysis of the C. elegans Argonaute family reveals that distinct Argonautes act sequentially during RNAi. Cell 127, 747–757 (2006).

12. Pak, J. & Fire, A. Distinct populations of primary and secondary effectors during RNAi in C. elegans. Science 315, 241–244 (2007).

13. Guang, S. et al. Small regulatory RNAs inhibit RNA polymerase II during the elongation phase of transcription. Nature 465, 1097–1101 (2010).

14. Guang, S. et al. An Argonaute Transports siRNAs from the Cytoplasm to the Nucleus. Sci. (New York, NY) 321, 537–541 (2008).

15. Félix, M.-A. & Braendle, C. The natural history of Caenorhabditis elegans. Curr. Biol. 20, R965–9 (2010).

16. Altuvia, S., Weinstein-Fischer, D., Zhang, A., Postow, L. & Storz, G. A small, stable RNA induced by oxidative stress: role as a pleiotropic regulator and antimutator. Cell 90, 43–53 (1997).

17. Liu, H. et al. Escherichia coli noncoding RNAs can affect gene expression and physiology of Caenorhabditis elegans. Nat. Commun. 3, 1073 (2012).

18. Tabara, H. et al. The rde-1 gene, RNA interference, and transposon silencing in C. elegans. Cell 99, 123–132 (1999).

19. Brooks, K. K., Liang, B. & Watts, J. L. The influence of bacterial diet on fat storage in C. elegans. PLoS One 4, e7545 (2009).

20. Melo, J. A. & Ruvkun, G. Inactivation of conserved C. elegans genes engages pathogen- and xenobiotic-associated defenses. Cell 149, 452–466 (2012).

21. Ashe, A. et al. piRNAs Can Trigger a Multigenerational Epigenetic Memory in the Germline of C. elegans. Cell 150, 88–99 (2012).

